# Where, who, and what counts under area-based conservation targets: A framework for identifying opportunities that benefit biodiversity, climate mitigation, and human communities

**DOI:** 10.1101/2023.03.24.534176

**Authors:** Brooke L Bateman, Emily Feng, Joanna Grand, Lotem Taylor, Joanna X Wu, Sarah P Saunders, Chad Wilsey

**Affiliations:** National Science Team, National Audubon Society, 225 Varick Street, New York, NY 10014, USA; Current address: Department of Ecology & Evolutionary Biology, University of Connecticut, Storrs, CT 06269-3043, USA; Current address: Department of Ecology and Evolutionary Biology, University of California, Los Angeles, CA 90095-7246, USA

**Keywords:** Climate change, biodiversity loss, human well-being, IPLC, 30 by 30, America the Beautiful, natural climate solutions

## Abstract

1. Area-based conservation targets, such as ‘30−30’, if strategically applied, can increase resiliency to climate change and provide co-benefits to people and biodiversity. However, protected areas historically were not designated within the context of global change, and human communities at highest risk are often overlooked in conservation planning.
2. To inform 30−30 conservation planning in the United States (i.e., America the Beautiful; ATB), we evaluated *where* US conservation opportunities exist by identifying habitats that can simultaneously benefit climate change mitigation and bird populations, as well as *who* lives in these areas and how conservation actions could both improve human well-being or potentially be at odds with local communities. To inform the equitable implementation of area-based conservation targets, we integrated maps of critical habitat for birds now and under a changing climate with carbon stocks and sinks and developed a prioritization framework to investigate the spatial alignment of these locations with areas identified as important for both human well-being and land-dependent human communities.
3. Although nearly 30% of US lands have some level of protection, only 6% of US lands (143 million acres) are managed for biodiversity and align with Bird and Carbon (BC) priorities, and <3% of protected US lands (59 million acres) align with priorities for Birds, Carbon, and Human well-being (BCH).
4. Of the 312 million acres of BCH priorities identified, 71% lack known protection or formal conservation plans (14% of US lands) and should be considered conservation opportunities that could simultaneously address the biodiversity and climate crises, and social inequities. Targeting these BCH areas for conservation action would contribute to more equitable benefits to marginalized communities, and could fulfill the ‘Justice 40’ commitment, which aims to allocate 40% of federal investments in climate benefits to marginalized communities (which, for the 30% goal under ATB equates to 12% of US lands).
5. At least 80% of all BCH priorities co-occur with Indigenous peoples and local communities (IPLCs) who have strong cultural and socioeconomic ties to the land, making it imperative to work with local communities to define *what counts* as conservation actions towards the 30% goal and what successful conservation outcomes that benefit biodiversity, climate change mitigation, and human communities look like.

## Introduction

Habitat loss and climate change are two of the most pervasive and detrimental threats to biodiversity globally (Grand et al., 2019; Jetz et al., 2007; Segan et al., 2016). Climate change also contributes to species declines (Baranov et al., 2020; Iknayan & Beissinger, 2020; Soroye et al., 2020; Stephens et al., 2016), as a consequence of range loss and/or exposure to climate change-related threats (Bateman, Taylor, et al., 2020; Bateman, Wilsey, et al., 2020; Bellard et al., 2012; Warren et al., 2018; Wilsey et al., 2019). Furthermore, conversion of natural ecosystems has accelerated climate change by contributing up to 20% of the world’s greenhouse gas (GHG) emissions through released carbon into the atmosphere (IPCC, 2018). This number is expected to increase as continued warming leads to accelerated carbon loss (IPCC, 2018). We are amid a dual climate change and biodiversity crisis globally and solutions are urgently needed. Global conservation area-based targets, such as the Kunming-Montreal Global Biodiversity Framework to protect 30% of lands and water by 2030 (hereafter ‘30−30’), and the Global Deal for Nature to protect 50% by 2050 (‘Half-Earth’), are focused on reversing biodiversity loss through the conservation and restoration of natural areas (Dinerstein et al., 2019). Recently, the US federal administration has committed to implementing 30−30 under the America the Beautiful (ATB) plan, which aims to improve “our economy, our health, and our well-being”, support Tribal priorities, and use science as a guide (*Conserving and Restoring ‘America the Beautiful,’* 2021). Most protected areas in the US have been established for geological or scenic preservation, with little focus on biodiversity or climate change (Aycrigg et al., 2013; Jenkins et al., 2015; Venter et al., 2014). While 27% of continental US lands (i.e. excluding Hawaii) are considered formally protected, bringing us close to the 30% target on paper, in practice, the majority of these lands are managed for multiple uses and ignore the Tribal Nations and biodiversity goals of ATB (Dreiss et al., 2022; Dreiss & Malcom, 2022). There is a growing consensus for the need to consider more than simply protection designation of pre-defined acreage targets, with a shift in focus to both biodiversity and carbon mitigation (carbon stores and sinks) in conservation planning. Identifying locations where such natural resources co-occur can guide conservation in a way that more effectively addresses the biodiversity and climate crises (Carroll & Ray, 2020; Dreiss & Malcom, 2022; Goldstein et al., 2020).

Of the few analyses that identify ‘optimal’ conservation portfolios that maximize biodiversity and carbon benefits simultaneously, many overlook the rights of people living in these areas, as well as the implications for displacing people and current land uses. Notably, area-based conservation targets, such as ATB, have the potential to perpetuate environmental injustices if consideration of human well-being and livelihoods are ignored (Menton et al., 2020). Movements such as Half-Earth have been criticized for solely focusing on conserving biodiversity and overlooking the human communities that would be affected by its implementation (Büscher et al., 2017; Napoletano & Clark, 2020; Schleicher et al., 2019). Given the interconnectedness of climate change, biodiversity loss, and human well-being, we must strive for a holistic solution with an emphasis on the human communities at highest risk from climate change and land degradation who have historically been marginalized, and even victimized, in conservation planning. Solutions should not only protect biodiversity and restore degraded ecosystems, but should also contribute to climate change stabilization and resilience more broadly (Arneth et al., 2020; Bateman, Wilsey, et al., 2020; Saunders et al., In Press; Secretariat of the Convention on Biological Diversity, 2020). Conserving natural habitats through the implementation of natural climate solutions (NCS) is a promising strategy for simultaneously addressing both the biodiversity and climate crises while providing ecosystem services (Fargione et al., 2018; Griscom et al., 2017). Restoring or maintaining biodiversity in natural ecosystems under the NCS framework can provide co-benefits to people, including clean and abundant drinking water from healthy watersheds; increased productivity from healthy soils; flood control from functioning wetlands; and temperature moderation from healthy forests (Drever et al., 2021; Fargione et al., 2018).

Moreover, access to natural areas in the US has been shown to have disparities across race, socioeconomic status, and housing composition, with non-white and low-income communities being grossly underrepresented (Landau et al., 2020; Wolch et al., 2014). Access to nature can influence overall human health and well-being and foster opportunities for green economic development (Romagosa et al., 2015; Wolch et al., 2014). Governing bodies have historically ignored social and environmental justice and disregarded how marginalized human communities (e.g., Indigenous groups and Black and brown communities) are included or engaged in setting large-scale conservation targets (Schleicher et al., 2019; Tallis & Lubchenco, 2014). In the US, the Biden administration has committed to ‘Justice 40’, an initiative which aims to allocate 40% of federal investments in climate and clean energy benefits to marginalized communities that are underserved and overburdened by environmental inequities. This commitment provides an opportunity to address these legacy injustices under ATB by ensuring 40% of the ATB area-based conservation target (which, for the 30% goal under ATB would equate to 12% of US lands) falls within areas of overlap among priority places for biodiversity, climate change mitigation, and marginalized communities. Actively incorporating marginalized and historically excluded communities (e.g., Indigenous Peoples and Local Communities [IPLC]; Melanidis & Hagerman 2022) into conservation planning, such as by understanding who lives in priority restoration areas and how they may be affected by land management decisions, is critical for reaching successful and equitable conservation outcomes (Fleischman et al., 2022). As conservation efforts have the potential to conflict with social and ethical values, we must work toward a new paradigm of ‘Just conservation’ that mitigates injustices (Vucetich et al., 2018).

Though nearly 30% of US lands are currently within the protected area network, it remains unclear how well these lands help us achieve goals for biodiversity and climate change, and to what extent they account for human well-being. In this context, protection designation and the question of what counts as “protected” should determine whether we have met (or how to meet) ATB targets that achieve multi-faceted goals for natural spaces. We must identify where conservation action provides maximum return on investment for biodiversity and climate change and who will benefit or be engaged in the process (Lamb & Schmidt, 2021). Here, we provide a prioritization framework for identifying the spatial distribution of areas that are important for biodiversity and climate change mitigation (the *where*) and the human communities whose well-being and land-dependent livelihoods could be impacted by conservation actions (the *who*), within the context of area-based conservation targets including ATB. We focus on birds as representatives of biodiversity, as they are monitored globally and can be effective indicators of overall biodiversity when conservation planning spans many species, habitats, and environmental gradients (Fraixedas et al., 2020; Lewandowski et al., 2010). To align with the spatial extent of ATB and available data, we conducted this work in the continental United States. The maps at the scale of the continental US can be used in combination with local datasets and in partnership with IPLCs to identify *what counts* toward the 30% goal under the ATB initiative and to design regional conservation plans addressing the needs of both biodiversity and people. This framework for integrating multiple priorities into a spatial assessment of area-based targets is one approach to achieving more ‘just conservation’ (Vucetich et al., 2018).

## Methods

### Framework

We identified priorities, or areas of opportunity to benefit biodiversity, climate mitigation, and human communities. Key terms and definitions for each are provided in Table 1. These refer to areas identified by our method as priorities, a definition commonly used in literature on spatial prioritization for biodiversity conservation(see, Belote et al., 2021; Dorji et al., 2018; Stralberg et al., 2018; Stralberg, Carroll, et al., 2020; all of which refer to areas identified by their methods as “priorities” or “priority areas”). These areas are a function of our analytical decisions and objectives, and thus do not represent the only priorities to be considered in 30 by 30 conservation planning. First, to identify areas that meet dual conservation targets for both birds and carbon (BC priorities), we mapped Existing NCS Areas, or areas of high carbon storage and active carbon sinks that align spatially with Climate Strongholds, which we defined as areas important for birds today and under future climate change scenarios. We defined these alignment areas as Bird and Carbon priority Areas to maintain, where protection or sustainable management and conservation actions should be implemented to protect irrecoverable carbon in areas of high value to birds (Figure 1). We also identified Potential NCS Areas, or areas with high potential to act as carbon sinks if anthropogenic disturbance is minimized, which align with Vulnerable Climate Strongholds, defined as Climate Strongholds at high risk of conversion. We refer to these areas of alignment as Bird and Carbon priorities to restore, where restoration has the highest potential to improve carbon storage, sequestration, and bird habitat (Figure 1). Together, these areas are our Bird and Carbon priorities (BC priorities). Next, we assessed the alignment of these BC priorities with spatial data on marginalized communities across the US to identify areas where conservation efforts could simultaneously benefit biodiversity, climate change, and equitable human well-being, and designated these areas Bird, Carbon, and Human priorities (BCH priorities; Figure 1). Finally, we assessed how these priorities intersect with IPLCs, including land-dependent livelihood and cultural importance factors (BC+IPLC priorities and BCH+IPLC priorities).

**Figure 1.**
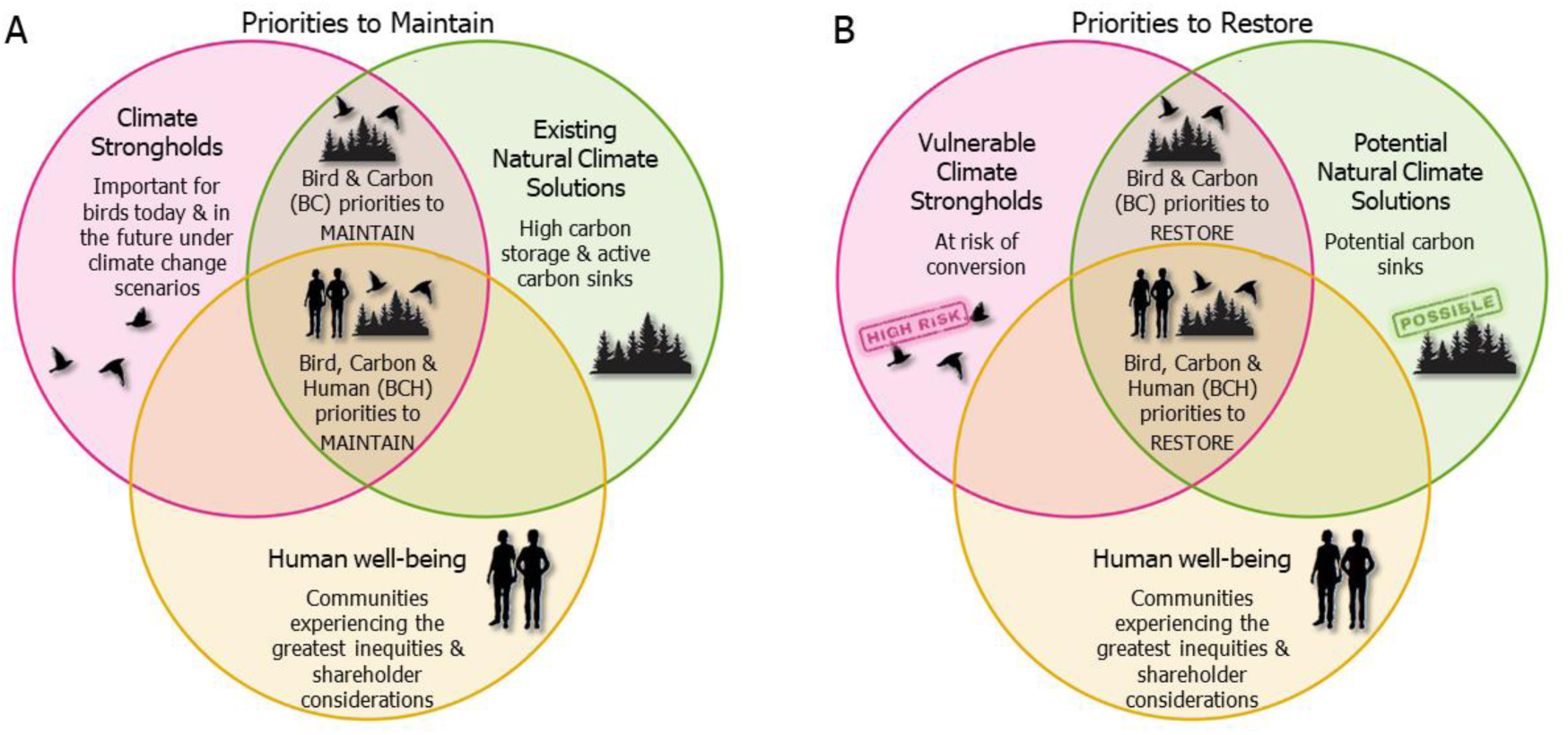
Conceptual diagram of how the Bird, Carbon, and Human priorities (BCH priorities), areas of alignment between bird climate strongholds, areas important for carbon storage and sequestration (natural climate solutions), and vulnerable human communities facing inequities (opportunities to equitably improve human well-being), are defined as, A) priorities to maintain and B) priorities to restore, based on the overlap of conservation priorities for birds (pink), carbon (green), and human well-being (yellow). Priorities to maintain are areas where Climate Strongholds and existing natural climate solutions are actively benefiting bird species resilience and carbon mitigation. Priorities to restore are areas where Vulnerable Climate Strongholds and potential natural climate solutions have potential benefits to bird resilience and carbon mitigation if restored. In each scenario (A and B), we focus on two areas of overlap 1) alignment of Climate Strongholds and natural climate solutions (benefits to Bird and Carbon – BC priorities) and 2) alignment of all three priorities (benefits to birds, carbon, and humans-BCH priorities). For both BC and BCH priorities to maintain and restore, we also consider areas with IPLCs considerations (i.e., BC+IPLC and BCH+IPLC) as additional priority area types (not shown).

**Table 1.** Terms with their acronyms and definitions that are used to describe conservation and restoration concepts that were developed, supported, and used in our analysis framework.

| Term | Acronyms and abbreviations | Definition | Data Description |
| --- | --- | --- | --- |
| Natural Climate Solutions | NCS | Management actions that mitigate greenhouse gas emissions and store and sequester carbon on working and natural lands (Fargione et al., 2018; Griscom et al., 2017) | See Existing and Potential NCS Areas. |
| Existing NCS Areas | Existing NCS | Areas of high carbon storage and active carbon sinks | Carbon storage = Total Ecosystem Carbon in 2015; carbon sequestration = Net Biome Productivity. (Liu et al., 2020) |
| Potential NCS Areas | Potential NCS | Areas with high potential to function as carbon sinks if anthropogenic disturbance is minimized | Carbon sequestration potential = Net Biome Productivity minus Net Ecosystem Productivity (Liu et al., 2020) |
| Climate Strongholds |  | Areas important for birds today and under future climate change scenarios | Top 40% of a prioritization-based landscape ranking of the continental U.S. based on current and future distributions of 557 bird species and habitat availability, while accounting for landscape condition, at a 1-km resolution. |
| Vulnerable Climate Strongholds |  | Climate Strongholds at high risk of conversion or with current high levels of conversion and degradation | Top 40% of a prioritization-based landscape ranking of the continental U.S. based on current and future distributions of 557 bird species and habitat availability, when landscape condition was not considered, at a 1-km resolution. |
| Bird and Carbon priorities to maintain | BC priorities to maintain | Areas where protection or sustainable management and conservation actions should be implemented to protect irrecoverable carbon in areas of high value to birds | Alignment of Climate Strongholds with Existing NCS areas |
| Bird and Carbon priorities to restore | BC priorities to restore | Areas where restoration has the highest potential to improve carbon storage, sequestration, and bird habitat | Alignment of Vulnerable Climate Strongholds with Potential NCS areas |
| Bird and Carbon priorities | BC priorities | Areas of alignment between bird climate strongholds and areas important for carbon storage and sequestration | Combined BC priorities to maintain and restore |
| Indigenous Peoples and Local Communities | IPLC | People within communities having the potential to be negatively affected by protected area designations and other conservation actions, including factors such as land-dependent livelihood and cultural importance factors human | Here, defined as Indigenous peoples, those with a high percentage of their population employed in the natural resources industry, or marginalized and low-income urban communities who are particularly prone to displacement due to green gentrification |
| Human Well-being |  | Areas of opportunities for establishing just and equitable outcomes for human communities | Here, defined as communities identified as socially vulnerable based on socio-economic considerations in addition to having at least one more inequity indicator in the following categories: climate change, pollution, health, and access to nature |
| Bird, Carbon, and Human Well-being priorities | BCH priorities | Areas where conservation efforts could simultaneously benefit biodiversity, climate change, and equitable human well-being (Can be divided into priorities to maintain and restore based on BC priorities to maintain and restore) | BC priorities with human-well-being considerations |
| Bird and Carbon priorities with IPLC considerations | BC+IPLC priorities | BC priorities that intersect with IPLCs, including land-dependent livelihood and cultural importance factors | BC priorities IPLCS considerations |
| Bird, Carbon, and Human Well-being priorities with IPLC considerations | BCH+IPLC priorities | BCH priorities that intersect with IPLCs, including land-dependent livelihood and cultural importance factors | BC priorities with both IPLC and human-well-being considerations |

### Bird priorities: Climate Strongholds

#### Data Description

We identified Climate Strongholds and Vulnerable Climate Strongholds for birds in 17 biogeographical groups (as per Taylor et al., 2022) using an optimization approach implemented with Zonation conservation planning software (Moilanen 2007). Birds were represented by previously published species distribution models developed by Bateman et al. (2020) at a 1-km resolution across North America for a baseline period (1981-2010) and for three future time periods (2020s, 2050s, and 2080s) under Representative Concentration Pathway (RCP) 8.5. For a list of species associated with each biogeographical group, please see S1 Data. The 17 biogeographical groups were based on habitat affiliations for 557 bird species (see S1 Data) and NCS focus areas (Bateman, Wilsey, et al., 2020; as per Fargione et al., 2018; Taylor et al., 2022) for seven ecosystems (forests, grasslands and rangelands, aridlands, interior wetlands, coastal wetlands, tundra, and urban and suburban systems) across four major regions of the continental US (eastern, central, western, and Alaska). For each biogeographical region (e.g. eastern forests), we calculated the spatial extent for each time period based on both current and future landcover projections that incorporated climate change, sea level rise, and human population growth (Alaska Center for Conservation Science, 2020; NOAA Office for Coastal Management, 2019; Rehfeldt et al., 2012; T. Sohl et al., 2016; T. L. Sohl, 2014; U.S. Environmental Protection Agency, 2017; U.S. Fish and Wildlife Service (USFWS), National Wetlands Inventory, 2020).

#### Data Analysis

We represented each species by its predicted suitability above a species-specific threshold (as per Bateman, Wilsey, et al., 2020; Taylor et al., 2022) and included projections from both breeding and nonbreeding seasons and all time periods under an ensemble of 15 GCMs as input features in Zonation. Following prior work to develop climate-informed bird prioritizations (e.g., Grand et al., 2019; Taylor et al., 2022), we used the Core Area Zonation algorithm to rank the landscape based on current and future climate suitability for birds, habitat availability, and landscape condition. For all ecosystems we used the Global Human Modification Index (Theobald et al., 2020) to represent degradation due to human activities. For forests, we also incorporated the Forest Landscape Integrity Index (Grantham et al., 2020) by selecting the maximum value between the two indices to identify the greatest disturbance threat. We used the inverse of the index to represent landscape condition (i.e., high degradation=poor condition). We limited condition layers to the extent of each biogeographical group for each time-period, and associated species layers with the corresponding condition layer from each time-period using Zonation’s grouping function to capture habitat quality and availability. We used Zonation’s ecological interactions function to downweight locations farther than a species’ natal dispersal capacity for each time period (Carroll et al., 2010; Grand et al., 2019; Moilanen et al., 2014; Rayfield et al., 2009), up-weighted at-risk species (derived from NABCI conservation status scores and climate vulnerability scores; Bateman et al. 2020; see S1 Data), and weighted present and near-future predictions higher than late-century predictions to address the increasing uncertainty of climate change projections over time (Knutti & Sedláček, 2013). To ensure that opportunities for conservation under ATB encompass ecologically distinct regions and are available across the continental U.S., we stratified rankings either by watershed boundaries (for coastal wetlands) or Bird Conservation Regions (for all other habitat groups; BCRs), the latter being distinct ecological regions with similar bird communities, habitats, and conservation issues (Bird Studies Canada and NABCI, 2014).

To identify Climate Strongholds, we selected the core (top 20%) and supporting biodiversity areas (top 40%-top 20%) of the landscape ranking (Jalkanen et al., 2020; Veloz et al., 2015) within each ecosystem (with the exception of (sub)urban where we used the core area (20%) due to the broad footprint of this system and generalist nature of species included). We selected he top 20 and 40% thresholds as they have been shown to be robust thresholds that capture more individuals per species than areas of lower priority (Jalkanen et al., 2020; Veloz et al., 2015). These top areas were predicted to have high climate suitability for birds and low human modification at present and/or under future climate change scenarios. To identify Vulnerable Climate Strongholds, we included both (1) areas not identified as Climate Strongholds but ranked in the top 40% of the prioritization when landscape condition was not considered (i.e., areas that have high climate suitability for birds but have been exposed to high amounts of human modification) and (2) areas in the top 40% of the difference between rankings with and without landscape condition (i.e., areas with the greatest potential to increase their value to birds if landscape condition was restored).

### Carbon Priorities: NCS Areas

#### Data Description

To identify existing and Potential NCS Areas, we identified locations with active (i.e., areas that currently function as significant carbon stores or sinks) and potential (i.e., areas that could have high carbon sequestration value if disturbance is minimized) carbon mitigation capacity. We used datasets from the Integrated Biosphere Simulator (IBIS) model to map carbon storage and flux (i.e., sequestration rate, or the annual amount of carbon exchange between an ecosystem and the atmosphere) in the contiguous US (Liu et al. 2020). To map areas of high carbon storage, we used Total Ecosystem Carbon (TEC) from the latest year available (2015), which measures aboveground and belowground carbon up to two meters. To map areas of high carbon sequestration, we used Net Biome Productivity (NBP), which measures the realized annual rate of carbon exchange, incorporating human and natural disturbances. To identify areas with high potential for increased carbon sequestration, we mapped annual Net Ecosystem Productivity (NEP), or carbon flux in the hypothetical absence of natural or human disturbance, and calculated the difference between NBP and NEP. We recognize this may overestimate the area with potential for increased carbon sequestration given some ecosystems require natural disturbances; however, data distinguishing natural and anthropogenic carbon losses were not available.

To map carbon storage and flux in Alaska, we used data from the Scenarios Network for Alaska and Arctic Planning (SNAP) program based on the Terrestrial Ecosystem Model simulations for Alaska and Northwestern Canada, averaged during 2001-2010 (Genet et al. 2015).

#### Data Analysis

We summarized flux rates across the lower 48 from 1980-2015 to obtain the average and maximum sequestration rate over time (measured in tons of carbon per year). We summarized high and low estimates of flux rates, calculated as the difference between carbon sequestration continuing at average historical rates from 1980–2015 (low) versus maximum historical rates (high). To calculate flux rates in Alaska, we subtracted heterotrophic respiration (HR) from vegetation Net Primary Productivity (NPP) to obtain NEP (net flux without accounting for loss due to disturbance (C/m^2^/yr). To calculate NBP (gC/m^2^/yr), we subtracted emissions due to disturbance (fire emissions from organic layer and vegetation burning) from NEP. Total carbon was calculated by summing the decadal average of soil and vegetation carbon pools (gC/m^2^).

We extracted carbon estimates within the biogeographical extent of the Climate Strongholds for each of the 17 biogeographical groups. We included all areas also mapped in the corresponding IBIS time series dataset of the dominant land cover class (i.e., forests, grasslands and rangelands, and aridlands) to ensure the carbon data aligned with the appropriate habitat. Existing NCS areas included areas in the top 20% of TEC for each ecosystem, or that had a positive flux based on NBP, indicating these areas currently have high carbon storage capacity or are actively sequestering more carbon than they emit. We chose the 20% threshold for carbon to be comparable, and for assessing potential spatial overlap, with our Climate Strongholds, based on best practices in prioritization assessments looking at multiple conservation values (Carroll & Ray, 2020; Morán-Ordóñez et al., 2017; Reside et al., 2017). Potential NCS Areas included areas where NEP was greater than NBP, indicating these areas could sequester more carbon if disturbance was minimized. We did not adjust carbon sequestration estimates for albedo, but acknowledge that in some ecosystems, especially high latitude conifer forests, reduced albedo can lead to a warming effect that offsets carbon storage benefits (Naudts et al. 2016).

### Bird and Carbon Priorities: The *Where*

Within each ecosystem, we identified Bird and Carbon priorities to maintain (BC priorities, maintain), where Climate Strongholds overlap with Existing NCS Areas (Figure 1). We considered these areas to have the greatest conservation benefit as they simultaneously help birds adapt to climate change while mitigating further carbon emissions. We also identified Bird and Carbon priorities to restore (BC priorities, restore) where Vulnerable Climate Strongholds aligned with Potential NCS Areas (Figure 1). In Alaska, we only calculated BC priorities to maintain due to lack of available carbon data incorporating human disturbance. We merged each ecosystem level BC priority layer across the US into a single layer of priorities to maintain and a single layer of priorities to restore. Finally, we overlaid BC maintain and restore priorities to create a single layer of BC priorities. As there was some overlap among habitats, any overlap between priorities to maintain and restore were considered priorities to maintain so as not to double count areas.

To evaluate how effectively these single (strongholds and NCS) and dual objective (BC) priorities captured bird communities, we developed potential bird richness layers based on modeled species distribution outputs (from Bateman, Wilsey, et al., 2020) by summing the climate suitability layers for both the baseline and future time periods for the suite of species in each biogeographic group. Each unique species-season combination was included as a separate input. We developed a future species refugia rank (e.g. species retention) layer by calculating the number of unique species-season layers retained for each pixel in both the present and the future, and then assessed retention for each stronghold, NCS, and BC priority (as per Taylor et al., 2022). Lastly, we assessed the current protected status within the different priorities within the context of US progress towards meeting ATB.

### Priorities for Human Well-Being: the *Who*

#### Data Description

We included five indicators of inequitable human well-being (hereafter “inequity indicators”) to identify where BC priorities aligned with opportunities for establishing just and equitable outcomes for human communities. We considered locations to have opportunities for improving equitable human well-being (hereafter “priorities for human well-being”) if communities were identified as socially vulnerable based on socio-economic considerations (Centers for Disease Control and Prevention 2018), in addition to having at least one more inequity indicator in the following categories: climate change, pollution, health, and access to nature. We obtained social data for human well-being from the Centers for Disease Control and Prevention’s (CDC) PLACES database (Centers for Disease Control and Prevention, 2021), the Council on Environmental Quality’s (CEQ) beta version of their Climate and Economic Justice Screening Tool (CEJST; U.S. Council on Environmental Quality, 2022), the EPAs environmental justice screening and mapping tool, EJScreen (US Environmental Protection Agency 2019), and the CDC’s Social Vulnerability Index (SVI; Centers for Disease Control and Prevention, 2018). Data from these sources come from a range of multi-year estimates (i.e., American Community Survey 2014-2018 and 2015-2019 estimates), analyses, and collection years that fall within an eight-year time window (2014-2021). In addition, health inequity indicators were based on 2019 Behavioral Risk Factor Surveillance System community surveys that assessed both the present status of individuals (i.e., physical inactivity and mental health as of 2019) and health history of individuals (e.g., history of heart disease and diabetes throughout the individual’s lifetime). All our human well-being analyses were calculated within 2010 US census tract units (hereafter, “communities”).

We used the methodology of the EPA and CEQ to highlight disadvantaged communities based on identifying census tracts that occur both 1) at or above the threshold for socioeconomic burden, and 2) at or above the threshold for one (or more) inequity indicators (U.S. Council on Environmental Quality, 2022). These thresholds were set by the EPA and the CEQ and are recommended to be used with these products, with the 80^th^ percentile nationally for an inequity indicator being considered a potential community burden, and 65^th^ percentile for socioeconomic burden. We used the social vulnerability index (SVI) to identify communities that are socioeconomically burdened and that are often more vulnerable and lack resources to prepare for and respond to catastrophic events. SVI considers socio-economic status, household composition and disability, minority status and language barriers, housing type and transportation availability. Communities above the 65^th^ national percentile for SVI were classified as socially vulnerable based on the CEQ threshold for socioeconomic burden (per CEJEST 2022). We considered a community disadvantaged if it occurs within a census tract that falls at or above the 65^th^ percentile of socio-economic vulnerability, and also is at or above the 80^th^ percentile as per the EPA and EJSCREEN (US Environmental Protection Agency 2019) for one or more inequity indicators (health, pollution, or climate). We applied the same threshold to the derived indicators we developed below.

We created a health inequity indicator, which considered the four health conditions (crude prevalence of asthma, coronary heart disease, diabetes, and low life expectancy derived from the National Center for Health Statistic’s US Small-Area Life Expectancy Estimates Project) which were included in CEJST and two additional health conditions from the PLACES database: the prevalence of physical inactivity and poor mental health. We added the latter two conditions because physical activity and mental health has been directly linked to access to greenery and outdoor spaces (James et al., 2015). If a community was ranked within the 80th percentile (as per EPA, see above) for any of these six health conditions, the tract was classified as having a health inequity.

To create a pollution inequity indicator, we combined several CEJST environmental burden indicators that focused on sources of pollution and non-clean infrastructure. These CEJST indicators were obtained from the Environmental Protection Agency’s (EPA) EJSCREEN environmental indicators (U.S. Environmental Protection Agency (EPA), 2019). Communities were classified as having this indicator if they were above the 80th percentile (as per EPA, see above) for any of the following conditions: wastewater discharge, proximity to hazardous waste facilities, Superfund National Priorities List (NPL) sites, Risk Management Plan (RMP) sites, diesel particulate matter (PM) exposure, traffic proximity and volume, or PM 2.5 in the air.

To create an indicator of climate change risk inequity, we kept the same indicators used to measure climate burdens in CEJST: a community was classified as having climate inequities if it was within the 80^th^ percentile (as per EPA, see above) for expected agricultural, building, or population losses due to natural hazards (Zuzak et al., 2021). For an access to nature indicator, we intersected census tracts with the US Protected Areas Database (PADUS version 2.1, U.S. Geological Survey, 2020) to identify communities without access to land formally protected for biodiversity (i.e., census tracts that did not intersect areas listed as GAP status 1-2 and Open Access in PADUS, U.S. Geological Survey, 2020).

#### Data Analysis

To identify BCH priorities, areas where priorities for birds, carbon, and human well-being overlapped, we intersected our priorities for human well-being and socially vulnerable communities across the US that had at least one additional inequity indicator (i.e., health, pollution, climate change risk, or access to nature), with our BC priorities.

### IPLC Considerations: The *What Counts*

#### Data Description

We focused on IPLCs, a subset of human communities having the potential to be negatively affected by protected area designations and other conservation actions, given these communities depend on the land for cultural preservation and livelihoods. Such communities include those of Indigenous peoples, those with a high percentage of their population employed in the natural resources industry, and marginalized and low-income urban communities who are particularly prone to displacement due to green gentrification. Green gentrification is a complex issue, but a large component of long-term displacement is increased housing costs associated with green infrastructure (Anguelovski et al., 2019). Despite the variation in land-dependencies existing within and between these three groups (e.g., economic living standards, subsistence living, health and wellness, culture, and spirituality), all of these dependencies are linked to overall human well-being and are reasons for mandating the freedom of choice and action in land management decisions (Burnette et al., 2018; McKinnon et al., 2016).

For our IPLC considerations, we used several data sources and retained our 80^th^ percentile threshold for consistency with our inequity indicators. For the Indigenous land indicator, we identified communities intersecting Indigenous land boundaries from the US Census Bureau’s 2019 National American Indian/Alaska Native/Native Hawaiian Areas (AIANNH) dataset (U.S. Census Bureau, 2019). These boundaries represent federally recognized American Indian reservations, off-reservation trust lands, state-recognized American Indian reservations, and state designated tribal areas. For the natural resources industry indicator, we included communities within the 80^th^ percentile for the percent of their working population in the natural resources industry from the US Census Bureau’s 2016-2020 American Community Survey (ACS) 5-Year Estimates of Selected Economic Characteristics (U.S. Census Bureau, 2020). Natural resources industries include agriculture, forestry, fishing, hunting, and mining. Finally, for the green gentrification indicator, we included communities within urbanized area boundaries classified by the 2010 US Census (U.S. Census Bureau, 2010) that were also in the 80^th^ percentile for the housing cost burden variable used in CEJST. Housing cost burden is the percent of households in each tract earning <80% of the Housing and Urban Development (HUD) Area Median Family Income by county (Office of Policy Development and Research (PD&R) Department of Housing & Urban Development (HUD), 2021) and are spending >30% of their income on housing costs.

#### Data Analysis

To identify priorities that may require IPLC engagement in conservation planning, we identified communities having at least one IPLC indicator (i.e., Indigenous land, natural resource industry, or green gentrification). We intersected these communities with our BC priorities and our BCH priorities and refer to these coincident areas as BC+IPLC priorities and BCH+IPLC priorities, respectively.

### Protection Status

To assess protected status for all priorities, we used GAP status, a measure of intent to conserve biodiversity classified as follows: GAP 1-2 are under permanent protection for conservation, GAP 3 is under protection but allows multiple uses and extraction, and GAP 4 has no known legal conservation mandate. Here, we classify GAP 1 and 2 lands as highly protected, given the mandate to protect biodiversity and natural land cover and excludes human resource extraction (Belote et al., 2021; Dreiss et al., 2022; USGS Gap Analysis Project, 2018). Gap 3 and 4 designated lands as they currently are managed may be inadequate to conserve lands in a way that maintains biodiversity, carbon mitigation values, and/or support human well-being, (Dreiss et al., 2022). However, Gap 3 and 4 lands have the potential to be beneficial to biodiversity, climate mitigation value, and human well-being with some adjustments, such as changes in management practices, restoration efforts, and/or increased access to nature (Dreiss & Malcom, 2022). We flattened the PADUS GAP status data layer (U.S. Geological Survey, 2020) across overlapping areas by retaining the highest level of protection for each area, following methods in Dreiss & Malcom (2022) and considered non-PADUS lands as GAP 4 since their protected status is unknown. Although we designate GAP 4 as “unprotected”, we acknowledge these areas may have local mandates or private land ownership and management that are not captured by PADUS. We calculated the percent of the total land area within the continental US, within sub-national regions determined by the US Global Change Research Program (National Climate Assessment Regions (NCA4) (USGCRP, 2017)), and within the identified areas for each of our priorities classified as GAP 1-3.

## Results

### Birds and Carbon Priorities: The *Where*

We identified 1.1 billion acres of BC priorities (BC priorities to maintain and restore, combined) across all ecosystems (Figure 2). BC priorities had equal mean potential species richness (49 species/1-km cell) as Climate Strongholds alone (49 species/1-km cell), but slightly higher than NCS areas alone (42 species/1-km cell) within the contemporary time-period (1981-2010). Climate Strongholds and BC priorities had slightly greater species retention (76% and 74%, respectively) than NCS areas (73%) under future climate change. Of the BC priorities across all ecosystems, priorities to maintain had slightly greater mean potential species richness (50 species/1-km cell) than priorities to restore (48 species/1-km cell) within the current time-period, and greater retention of species under future climate change (76% and 74%, respectively).

**Figure 2.**
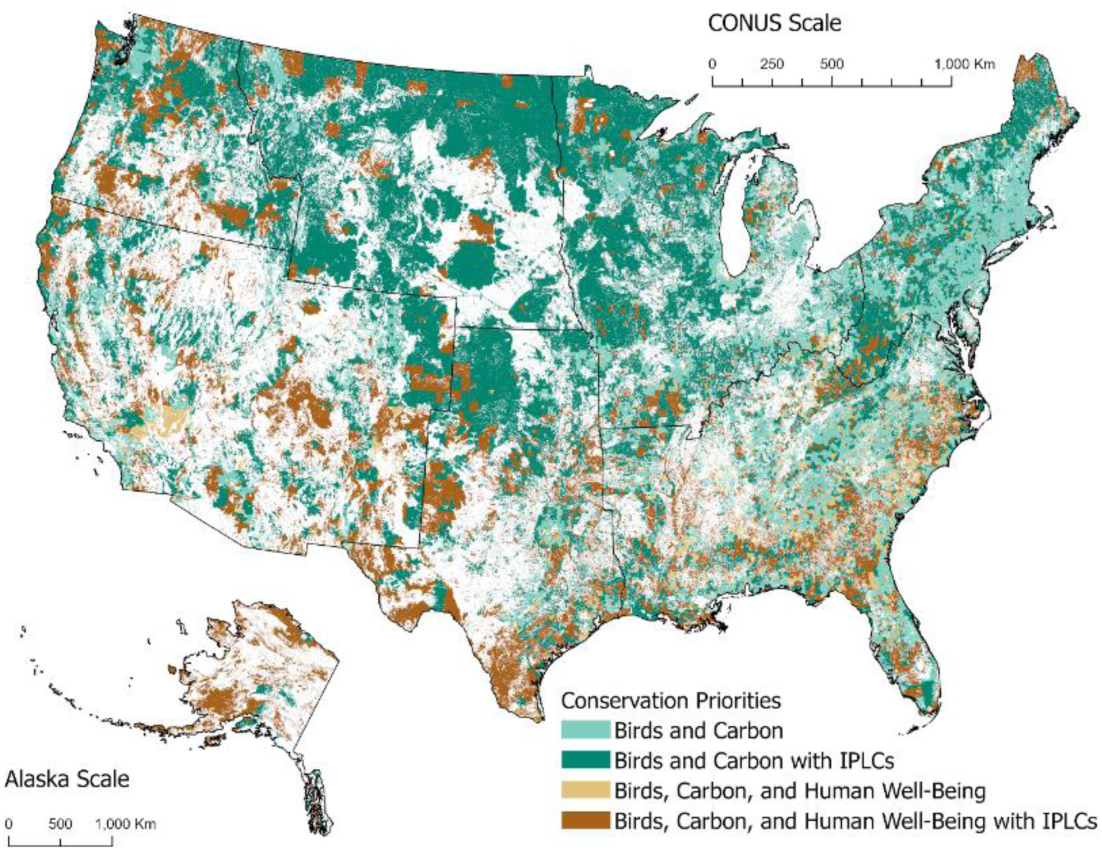
US lands identified with the highest potential for birds and carbon (BC priorities), as well as potential to promote equitable human well-being (BCH priorities) if maintained or restored. Darker color tones indicate conservation priorities occurring in communities that rely on the land for cultural preservation and/or socioeconomic factors (BC+IPLC and BCH+IPLC).

BC priorities store a total of 103.0 billion tons of carbon, and, over time, these priorities have functioned as net carbon sinks, sequestering more carbon than they emit (net 139 million tons carbon per year). BC priorities to Maintain sequester, on average, 106.8 million tons of carbon per year, and have a potential sequestration of up to 146.3 million tons of carbon per year. Similarly, BC priorities to Restore sequester, on average, 32.6 million tons of carbon per year and have a potential sequestration of up to 71.0 million tons of carbon per year if disturbance is reduced.

A total of 74% of BC priorities occur on unprotected land with no conservation mandate (see S3 Table). These are potential opportunities under 30−30 to conserve biodiversity and mitigate carbon and comprise 43% (980 million acres) of the US (Table 2, Figure 3, S4 Table). Currently 13% (305 million acres) of the continental US is under strict protection (GAP 1-2), with an additional 18% (410 million acres) under multiple-use conservation protection (GAP 3, Table 2. Thus, most of the US (69%) does not have any conservation protection. Nearly 50% of current GAP 1-2 protected lands also coincide with BC priorities (Table 2, Figure 3), which equates to 6% of US lands (143 million acres) that could count towards the ATB target (Table 2, Figure 3, S4 Table). Currently, 48% of GAP 3 managed lands overlap with BC priorities, equating to 9% of US lands (200 million acres) that are potential opportunity areas for increased protection levels to count towards 30−30 while benefiting birds and carbon (Table 2, Figure 3, S4 Table).

**Figure 3.**
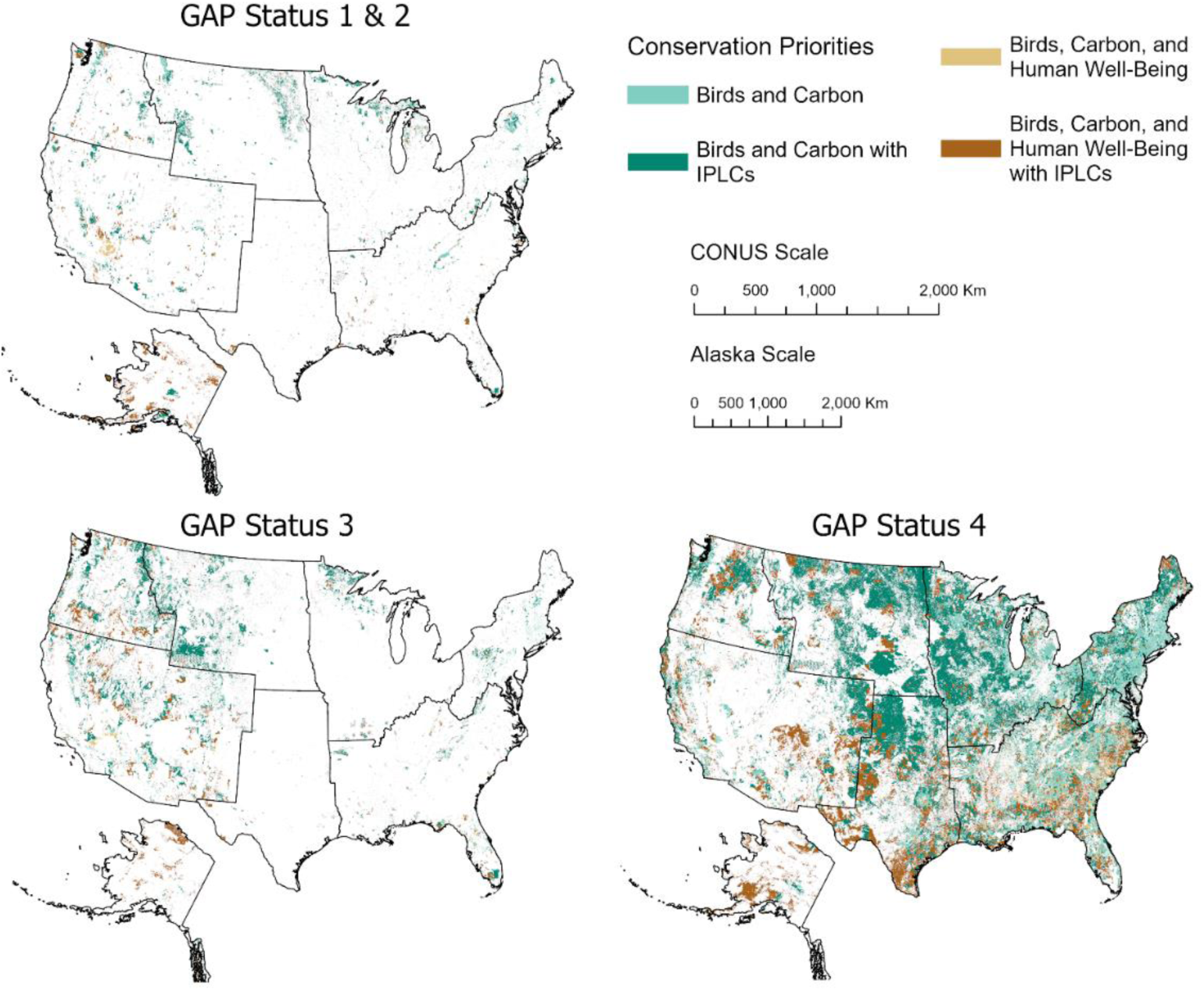
US lands identified with the highest potential for birds and carbon (BC priorities), as well as potential to promote equitable human well-being (BCH priorities) if maintained or restored according to their current GAP status. Darker color tones indicate conservation priorities occurring in communities that rely on the land for cultural preservation and/or socioeconomic factors (BC+IPLC and BCH+IPLC).

**Table 2.** Percentages of land area relative to the total area within each region that are classified as different priorities under each protection status, with Bird and Carbon (BC) priorities in green columns and Bird, Carbon, and Human (BCH) priorities in tan columns. Darker color tones represent the areas within those BC and BCH priorities that have land-dependent Indigenous Peoples and Local Communities (IPLCs).

| All Inland Area |  |  |  | BC Priorities |  |  | BCH Priorities |  |  | BC Priorities with IPLCs |  |  | BCH Priorities with IPLCs |  |  |
| --- | --- | --- | --- | --- | --- | --- | --- | --- | --- | --- | --- | --- | --- | --- | --- |
| Region | GAP 1-2 | GAP 3 | GAP 4 | GAP 1-2 | GAP 3 | GAP 4 | GAP 1-2 | GAP 3 | GAP 4 | GAP 1-2 | GAP 3 | GAP 4 | GAP 1-2 | GAP 3 | GAP 4 |
| US | 13.26 | 17.85 | 68.89 | 6.20 | 8.68 | 42.69 | 2.55 | 2.97 | 13.56 | 5.12 | 7.67 | 33.24 | 2.36 | 2.78 | 11.71 |
| Alaska | 6.36 | 3.54 | 90.10 | 2.01 | 1.16 | 2.70 | 1.51 | 0.98 | 2.30 | 1.81 | 1.14 | 2.62 | 1.51 | 0.98 | 2.30 |
| Southwest | 2.69 | 7.05 | 90.26 | 1.16 | 2.78 | 4.68 | 0.47 | 1.03 | 2.07 | 0.93 | 2.45 | 3.92 | 0.36 | 0.92 | 1.92 |
| Northern Great Plains | 1.19 | 2.57 | 96.24 | 0.86 | 1.66 | 5.85 | 0.03 | 0.08 | 0.70 | 0.80 | 1.59 | 5.77 | 0.03 | 0.08 | 0.69 |
| Northwest | 0.92 | 2.88 | 96.20 | 0.53 | 1.58 | 2.21 | 0.16 | 0.54 | 0.77 | 0.47 | 1.46 | 1.95 | 0.16 | 0.54 | 0.75 |
| Midwest | 0.81 | 0.49 | 98.70 | 0.62 | 0.43 | 8.02 | 0.08 | 0.05 | 0.86 | 0.48 | 0.39 | 5.98 | 0.06 | 0.04 | 0.64 |
| Southeast | 0.71 | 0.76 | 98.53 | 0.55 | 0.63 | 8.63 | 0.20 | 0.22 | 3.64 | 0.33 | 0.37 | 4.78 | 0.16 | 0.15 | 2.45 |
| Northeast | 0.40 | 0.41 | 99.19 | 0.36 | 0.36 | 4.09 | 0.03 | 0.03 | 0.52 | 0.20 | 0.20 | 2.28 | 0.02 | 0.02 | 0.40 |
| Southern Great Plains | 0.18 | 0.15 | 99.67 | 0.11 | 0.08 | 6.50 | 0.06 | 0.05 | 2.70 | 0.09 | 0.07 | 5.94 | 0.06 | 0.05 | 2.56 |

### Priorities for Human Well-Being: the *Who*

A total of 19% of US lands (438 million acres) align with BC priorities and are also within communities identified as priorities for human well-being (BCH priorities; Table 2, Figure 2). A quarter of the US population lives in communities with these BCH priorities (hereafter “BCH communities”). Less than 3% of US lands are BCH priorities that are currently protected (Table 2). Of these BCH priorities, 71% are unprotected and an additional 15% are within GAP 3 designation, which amounts to 380 million acres of conservation opportunity across the continental US (or 14% and 3% of the US, respectively; Table 2).

Of the community inequity indicators considered in this analysis, BCH communities frequently had less access to nature (91% of the population in BCH communities), followed by chronic health conditions (73%), pollution exposure (62%), and high exposure to climate change risk in the continental US (47%) (Figure 4b). Overall, 95% of populations in BCH communities face multiple inequities in a given location (Figure 5). Although the combination of inequalities, dominant inequities, and magnitude of overlap varied regionally (Figure 4b, Figure 5), high community inequity was prominent in the Southern Great Plains, Northern Great Plains, and Southeast.

**Figure 4.**
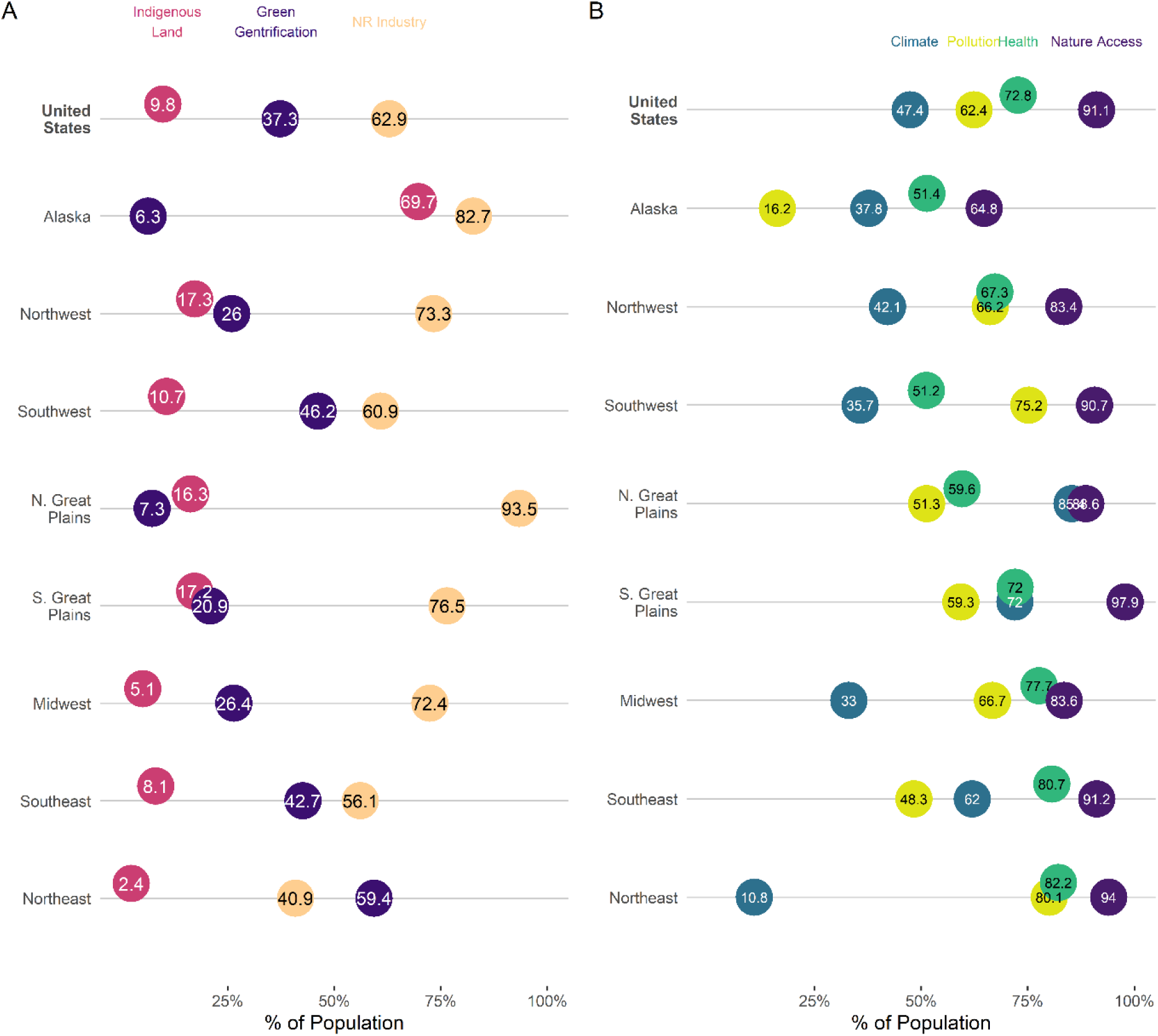
A) The percent prevalence of indigenous peoples’ and local communities’ (IPLC) considerations (green gentrification potential, indigenous land, or natural resource [NR] industry occupations) within populations of land-dependent communities that overlap Bird and Carbon priorities. B) The percent prevalence of human inequities (climate, health, pollution, or nature access) within populations of socially vulnerable communities that overlap Bird and Carbon conservation priorities (priorities for birds, carbon, and human well-being [BCH]). Percentages are relative to the estimated populations summed across census tracts within each US region.

**Figure 5.**
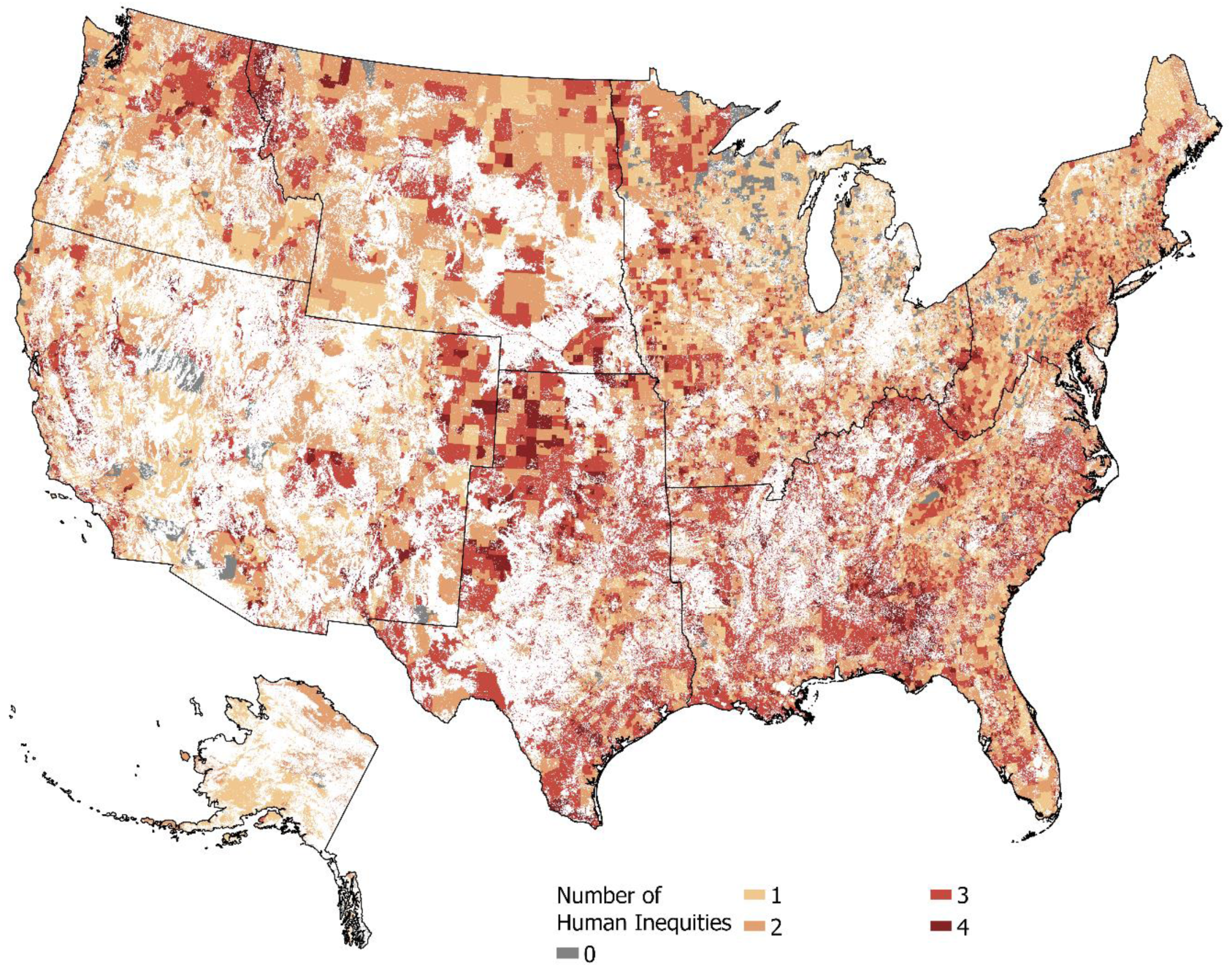
Bird and Carbon (BC) conservation priorities overlap with human communities experiencing co-occuring inequities. This map shows the number of co-occurring inequity indicators (including climate, pollution, health, and nature access) experienced by the human communities overlapping BC priority areas. Geographic region boundaries are shown in black line.

### IPLC Considerations: The *What Counts*

Across all communities with BC Priorities, IPLC considerations affected a greater percentage of the population (36%) than community inequities (30%). Of the 1.32 billion acres of BC priorities in the US, regardless of protection status, 80% also have indicators of IPLC considerations (i.e., BC+IPLC priorities) and 29% are in communities with inequitable human well-being in addition to having IPLC considerations (i.e., BCH+IPLC priorities; S3 Table). A total of 72% BC+IPLC priorities and 69% of BCH+IPLC priorities are unprotected, with a further 17% and 16% in GAP 3 lands, respectively (S4 Table). Across the continental US, 43% of GAP 3 lands coincide with BC+IPLC areas and 15% coincide with BCH+IPLC areas (the percentage of GAP3 in these priorities out of the total US GAP3 area, Table 2, S4 Table).

Of the population living within BC priorities and in communities with IPLCs, the natural resource industry indicator was the most prevalent IPLC consideration (63%) across the continental US. Natural resource industry dependence was highest in central and western regions, and lowest in the east (Figure 4a, S5 Table). The green gentrification indicator was the next most prevalent, affecting 37% of the population across the US, but percentages were greatest in the Northeast (59% of population) Southwest (46% of population) and the Southeast (43%; Figure 4a, S5 Table). Ten percent of this US population (BC + IPLC, and up to 70% in Alaska) resides in communities that share a mix of legal boundaries belonging to tribal and non-tribal governments, and state and federally recognized American Indian lands. When evaluating priority areas to maintain and restore across all regions, populations with high potential for green gentrification were more prevalent in priorities to restore while populations associated with Indigenous land and natural resource industry were higher in priority areas to maintain (S5 Table).

## Discussion

Currently, of the 13% of US lands with a legal mandate for biodiversity protection, only 6% align with BC priorities and <3% overlap with BCH priorities, revealing that we have not been successful to date in incorporating these priorities within our current protected area portfolio. Here, we identified an additional 980 million acres (43% of US lands) of unprotected land as potential opportunities to conserve BC priorities, and nearly 312 million acres (14% of US lands) representing opportunities to conserve BCH priorities. Focusing conservation efforts on these lands would meet the ATB goals and help address the dual climate and biodiversity crises (Carroll & Ray, 2020; Stralberg, Arseneault, et al., 2020), while simultaneously benefitting human communities.

Meeting ATB targets by increasing or adding protections to existing multiple-use areas (GAP 3) covering nearly 18% of US lands (e.g. Dreiss & Malcom, 2022) would not address climate change nor human equity. We found that 49% of GAP 3 lands have high value for birds and carbon (totaling 9% of US lands). In addition, we found only 19% of US lands with conservation protection, and 17% of multiple-use lands, also align with BCH priorities (each covering just under 3% of US lands). Protected areas historically have not been designed with biodiversity, climate change or equitable human well-being in mind. More so, conservation practices have been focused on optimizing resource extraction or preserving recreational, scenic, or geological value on land that has been colonized (Aycrigg et al., 2013; Jenkins et al., 2015; Rudd et al., 2021; Venter et al., 2014). These same practices have also led to restricted access to protected lands and lack of inclusion in conservation decision making, with the brunt of the inequity falling on communities of color, rural communities, and IPLCs (Rudd et al., 2021). Therefore, we must look beyond the current protected lands system to fulfill our commitment under ATB.

Furthermore, the Biden administration’s Justice 40 initiative funding target, which aims to allocate 40% of climate change funding to marginalized communities, could be integrated into ATB, for example by ensuring that at least 40% of the ATB area-based conservation target, or 12% of the US (i.e. 40% of the 30% area based target), is allocated for conservation or restoration action within vulnerable communities. Here, we identified nearly 14% of US land (currently unprotected) where conservation could improve equitable human well-being alongside Bird and Carbon priorities. Our results indicate 95% of socially vulnerable populations located in BC priorities face multiple inequities, including 91% without adequate access to nature, and >70% with chronic health conditions, which is consistent with other work demonstrating that communities of color in the US live in areas with more human modification (Landau et al., 2020) and less access to nature (Wolch et al., 2014). Given that access to green spaces promotes increased physical activity, and thus improves human health and well-being (Romagosa et al., 2015; Wolch et al., 2014), prioritizing conservation efforts within these highly modified habitats would likely benefit human well-being in addition to biodiversity and climate mitigation. A special focus could be given to regions with relatively high community inequity, including areas in the Southern Great Plains, Northern Great Plains, and Southeast (Figure 5).

Conservation actions in these priority areas will necessitate working closely with IPLCs to develop conservation solutions that support their well-being and land dependency needs and count towards ATB. We found 80% of BC priorities fall within IPLCs (BC+IPLC priorities), and most of these lands are not currently protected. The high overlap of these areas is key for addressing biodiversity loss and climate change within IPLCs, and highlights the need to explore localized conservation actions that account for people’s cultural values and livelihoods.

We considered three IPLC attributes that have been historically affected by conservation and land protection decisions, acknowledging that several more IPLC attributes could be considered and thus, our estimates of conservation areas in need of IPLC actions are likely an underestimate. ATB has set a target to inclusively conserve and restore, and explicitly recognizes many human uses of lands and waters can benefit natural systems including working lands (*Conserving and Restoring ‘America the Beautiful,’* 2021). This is an important framing, as even with expansion of protected areas, biodiversity loss and extinction rates continue to remain high (Crist et al., 2021). Therefore, protection status alone does not guarantee conservation success without additional management actions (Michel et al., 2021; Wauchope et al., 2022; Wood et al., 2014). This may be the case within the context of climate change, where proactive management strategies need to be implemented to continuously adapt to changing environmental conditions (Schuurman et al., 2020; Wu et al., 2018, 2022). As part of this approach, we must identify what counts under ATB and engage private landowners in conservation and stewardship of the land in a manner that can sustain both nature and the people that depend on it (Naugle et al., 2020). This includes working closely with rural and working lands communities, as 63% of our IPLCs within BC priorities (77% when only considering IPLCs with BC priorities to maintain, S5 Table) are employed in the natural resource industry across all regions and priorities in our study. Working lands, if managed appropriately, can be beneficial for both carbon sequestration (Fargione et al., 2018; Griscom et al., 2017) and biodiversity conservation (Brockington et al., 2018). For example, ranchers can implement grazing practices that are bird-friendly, thereby promoting resilient grassland bird communities (Michel et al., 2020). These strategies will be particularly critical in the Northern and Southern Great Plains, Alaska, Northwest, and Midwest where communities with BC priorities have high dependence on natural resources for livelihoods.

Two aspects of the IPLC analysis merit further discussion. First, Indigenous lands are home to 10% of the US population within IPLCs with BC priorities, but this proportion was as high as 70% in Alaska. Given dispossession and forced migration, Indigenous lands have been reduced by nearly 99% of their historic extent (Farrell et al., 2021). However, indigenous-managed lands have been shown to have greater vertebrate species richness and threatened species richness than traditional protected areas. This, highlights the importance of Indigenous land management, stewardship, and traditional knowledge to conservation (Garnett et al., 2018; Schuster et al., 2019; West et al., 2006). Focusing on collaborations supporting and enhancing Indigenous land management or community-led protected area establishment may offer a path towards reaching ATB conservation targets that benefits human well-being and elevates Indigenous land rights. The human impact of conservation decision-making will also be strong in Priority Areas to Maintain, as we found both natural resource industries and Indigenous lands IPLC populations were higher within these areas. Given the legacy of displacement and protection of land without collaboration within areas of high conservation value such as these, key conservation opportunities should be identified in partnership with local IPLCs. Private lands are crucial for both sustaining biodiversity and climate change mitigation, and thus local and co-produced conservation solutions that move beyond singularly relying on traditional protected area mandates, will be critical going forward (Brockington et al., 2018; Lamb & Schmidt, 2021; Naugle et al., 2020).

Second, about 37% of BC+IPLC communities across all regions were at risk of green gentrification, especially within the south and eastern US. This risk was notably higher within priority areas to restore across all regions. Conservation practitioners must acknowledge that restoration projects can both improve human well-being and have unintended consequences for local communities. Green gentrification warrants particular attention given the likelihood of human displacement associated with restoration or wilding efforts in urban settings (Anguelovski et al., 2019). New urban green areas have been associated with declines in populations of people of color, indicating that in some cases the co-benefits of ecosystem restoration or natural climate solutions resilience projects may cause unintended consequences for vulnerable communities (Shokry et al., 2020). The risk of displacement may be highest in areas vulnerable to climate change (e.g. high sea-level rise risk areas), especially close to downtown locations and near active transportation lines (Rigolon & Németh, 2020; Shokry et al., 2020). The focus should be on identifying where urban neighborhoods align with areas for potential conservation and resiliency projects, and working with local communities to help identify where and how to undertake such efforts (Anguelovski et al., 2019). This includes identifying and implementing anti-displacement tools (e.g., community land trusts) to secure affordable housing in the vicinity of conservation projects and ensuring that conservation projects are desired by the local community prior to their commencement (Anguelovski et al., 2019; Shokry et al., 2020).

Together, these BC priorities, through actions to maintain, manage, and restore under ATB, have the potential to sequester an additional 536.3 million tons of carbon dioxide per year, which equates to 23.2% of the 2016 Paris Agreement commitment (Bateman et al., 2021). Additionally, we provide a spatial roadmap on *who* conservation efforts should benefit, by integrating the Justice 40 commitment within ATB and identifying triple-priority areas that could simultaneously contribute to achieving goals for biodiversity, climate change mitigation, and historically marginalized peoples. Lastly, we highlight the importance of engaging IPLCs in conservation, and that defining *what* counts as conserved under ATB must be done in collaboration with local communities. To stabilize climate change, reverse biodiversity loss, and improve human quality of life simultaneously, conservation actions must shift away from historical practices that have excluded Indigenous communities for the sake of “protection” and perpetuated green gentrification that marginalizes people with low socioeconomic status (largely Black and brown populations).

We present a prioritization framework that aims to couple ecological values with human well-being. While our findings and conclusions offer a quantitative assessment of the ongoing challenges facing co-beneficial conservation planning and environmental equity, we acknowledge that conservation planning inherently requires many subjective decisions, values, and objectives. Variation in input values or threshold choice can yield different priority areas. While we recognize this introduces bias, an assessment of that bias was not our goal, and several other studies have addressed the effect of different weighting, thresholds, and valuation schemes in conservation assessments (Belote et al., 2021; Carroll et al., 2017; Karimi et al., 2017; Taylor et al., 2022; Whitehead et al., 2014). In addition, we used commonly applied and recommended thresholds for the input values we used (see methods).

We also recognize there are limitations to, and implications of, the datasets used in this study. For our measures of NCS areas we acknowledge that in some locations carbon sequestration rates may be overestimated due to albedo or use of NEP estimates. However, only a small amount of our study area included high latitude conifer forest, minimizing the implications of albedo, and though the use of NEP may slightly overestimate, it still allows for an estimation of relative sequestration potential of a location. We excluded areas of agricultural production for cropland to account for the need for food production, but recognize that some of these areas could have high carbon value (Mehrabi et al., 2018). The social datasets used in this analysis vary in their temporal resolution and the years in which they were collected. For instance, several of the health inequity indicators are based on the lifelong medical histories of individuals. Since health and other socio-economic variables in specific locations change rapidly over time, either through changes to individuals or their movement between communities, this also introduces uncertainty into our results which aim to capture current conditions in an area. Conservation management decisions also take time to plan and implement which further exacerbates the temporal uncertainty of results based on these temporally explicit data.

An additional level of spatial data uncertainty is introduced when using Census tract data. Census tracts represent a unit of area of homogeneous and consistent population size, and therefore vary in size and shape across the US, but have been shown to be highly correlated with more spatially explicit data especially within areas of larger population size (Kramer et al., 2010).y. However, census tracts have been readily used in looking at the intersection between conservation and human data, including measuring access to nature (Landau et al., 2020; Spotswood et al., 2021). We should note that access to nature as measured by overlap (intersection) with public lands formally protected for biodiversity, is only one way to quantify nature access, as is our metric for green gentrification. Other metrics, such as walking distance to green spaces and/or the quality and quantity of green spaces within a community, including size and location of green spaces and proximity to transportation (Rigolon & Németh, 2020) should be evaluated in future research.

Lastly, we selected a small subset of possible IPLCs that are nature dependent (Fedele et al., 2021) and could be affected by conservation actions. Conservation practitioners should also consider other local groups or Indigenous rights (e.g., Ojibwe treaty rights to hunt, fish, and manage wildlife and plant resources in the Great Lakes region https://glifwc.org/), or local Environmental Justice groups) that may not be represented in census data.

Our approach exemplifies only one potential avenue for the strategic implementation of ATB. Other approaches may consider terrestrial biodiversity as a whole, climate corridors, climate velocity metrics (e.g., Carroll et al. 2017), or different social values (e.g., natural capital assets, cultural services; Bryan et al. 2010) for example. Thus, we offer our analysis framework as a starting point for the continued development of a socio-environmental approach to conservation planning designed to address the most pressing human and environmental crises of our time.

### Conclusions and Recommendations

Conservation practitioners should help support communities in sustainable management of local natural spaces (Fraser et al., 2006). Programs that incentivize maintaining natural land cover or provide guidance on extractive practices that minimize biodiversity and carbon loss, will be key within priority biodiversity and carbon areas. For example, in the western US, regenerative grazing programs enable ranchers to keep their lands working over the long-term while incentivizing the use of bird- and climate-friendly management practices. In the northeast, maple syrup producers are incentivized to improve songbird habitat on their land by creating a market for products harvested on land managed for birds that can also provide carbon benefits. Creating tools for landowners to quantify the biodiversity, ecosystem resilience, and climate change mitigation value of their properties will help to support market-based conservation solutions that can keep ecosystems intact for the foreseeable future (Bateman et al., 2021; Fargione et al., 2018). In addition to market-based solutions, identifying conservation solutions that also provide community and human health and well-being benefits can help achieve both restoration and nature access goals.

Ongoing and future conservation within the current global context must be strategic, science- and Indigenous-led, and ambitious, but also intentional and inclusive, ensuring local communities and conservation practitioners work together towards mutual goals. Here, we provide a framework for how we can integrate conservation and human community values within the context of area-based conservation to move the US towards more ‘Just conservation’ (Vucetich et al., 2018). To stabilize and reverse the biodiversity and climate crises, places of high value for birds that align with ecosystem carbon storage and sequestration are recommended as high priority for conservation actions. Yet, we recognize that local communities may be at odds with these identified priorities. In such cases, conservation planning should account for the needs of the community and the desired conservation outcomes (Fraser et al., 2006; Redpath et al., 2013). As progress is made towards 30 by 30 goals, the best conservation plans will promote human well-being alongside actions that protect and conserve nature and help stabilize our climate.

## Acknowledgments

This work was funded by the John D. and Catherine T. MacArthur Foundation G-1511-150388. We thank Geoff LeBaron, National Audubon Society, and Audubon state office staff for reviewing bird species lists. We thank Pedro Hernandez for early conversations on how we can integrate Justice 40 goals into ATB, and Robert Harris, National Audubon Society, for reviewing and providing comments on the manuscript with an EDIB lens. We thank Benjamin M Sleeter, Western Geographic Science Center, USGS, for contributions to and support of the carbon science components.

## Supplemental Information

S1 Data. Climate Stronghold Species Lists with Weights

S2 Data. Species Ecosystem Geography List

S3 Table. Percentages of land area for different priorities under each current protection status relative to the entire Bird and Carbon priority area contained within each US region. Bird and Carbon (BC) priorities are in green columns and Bird, Carbon, and Human (BCH) priorities in tan columns. Darker color tones represent the areas within those BC and BCH priorities that are within land-dependent Indigenous Peoples and Local Communities (IPLCs).

S4 Table. Regional acres of land area in the US within different Bird and Carbon (BC) priorities and their current protection status, with BC priorities in green columns and Bird, Carbon, and Human (BCH) priorities in tan columns. Darker color tones represent the areas within those BC and BCH priorities that are within land-dependent Indigenous Peoples and Local Communities (IPLCs).

S5 Table. Percentages of three cultural and livelihood land dependencies that could be impacted by conservation decisions (high rates of natural resource industry employment, presence of indigenous land, and green gentrification potential) that make up the total estimated population size in census tract communities within each U.S. region that have both Bird and Carbon (BC) priorities and Indigenous Peoples and Local Communities (IPLCs). These rates are compared for IPLCs with only BC priorities to maintain (Maintain), only priorities restore (Restore), and all IPLCs that overlap the combined maintain and restore BC Priority area (Both). The Maintain-Restore Difference column shows whether the percent population impacted by each land dependency was higher (and by how much of a percent) in those tracts with BC priorities to maintain or those in need of restoration.

