## Supplementary figures and images for "Where, who, and what counts under area-based conservation targets: A framework for identifying opportunities that benefit biodiversity, climate mitigation, and human communities"

### S3 Table

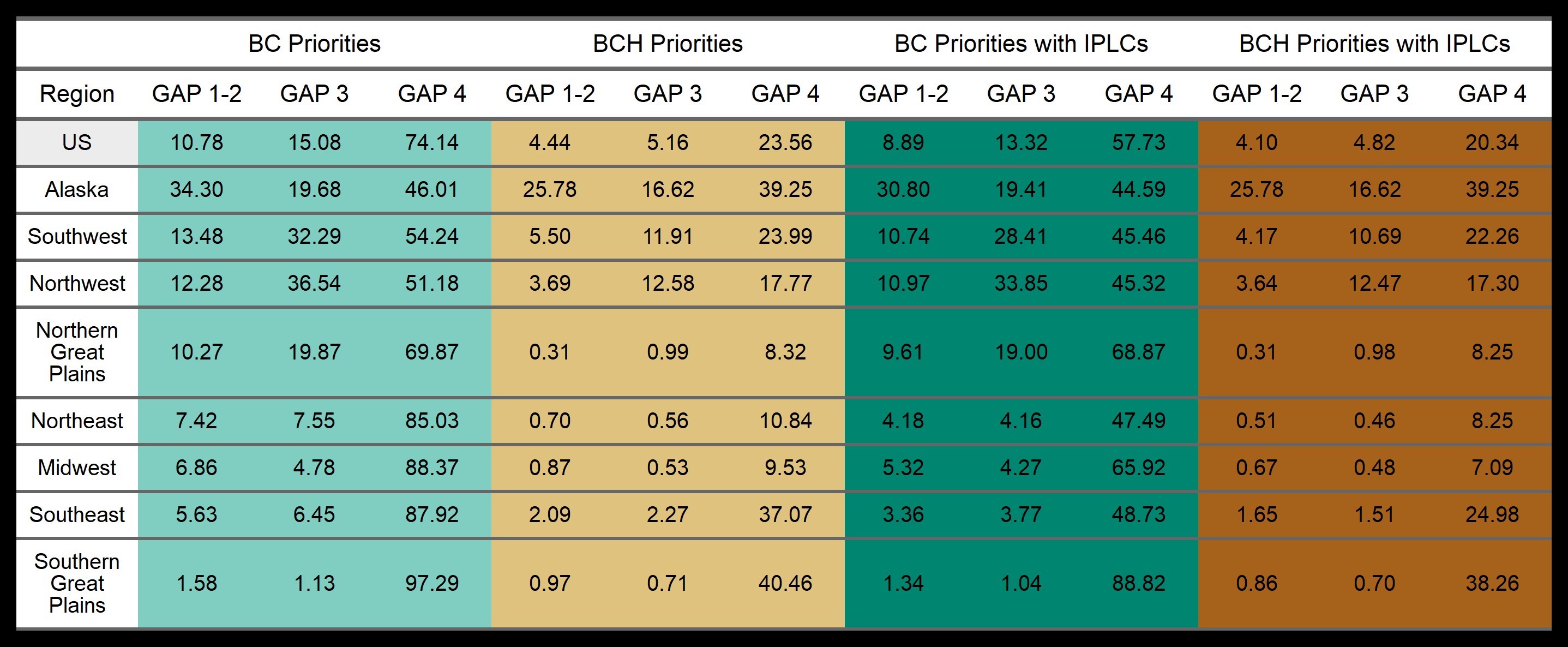

### S4 Table

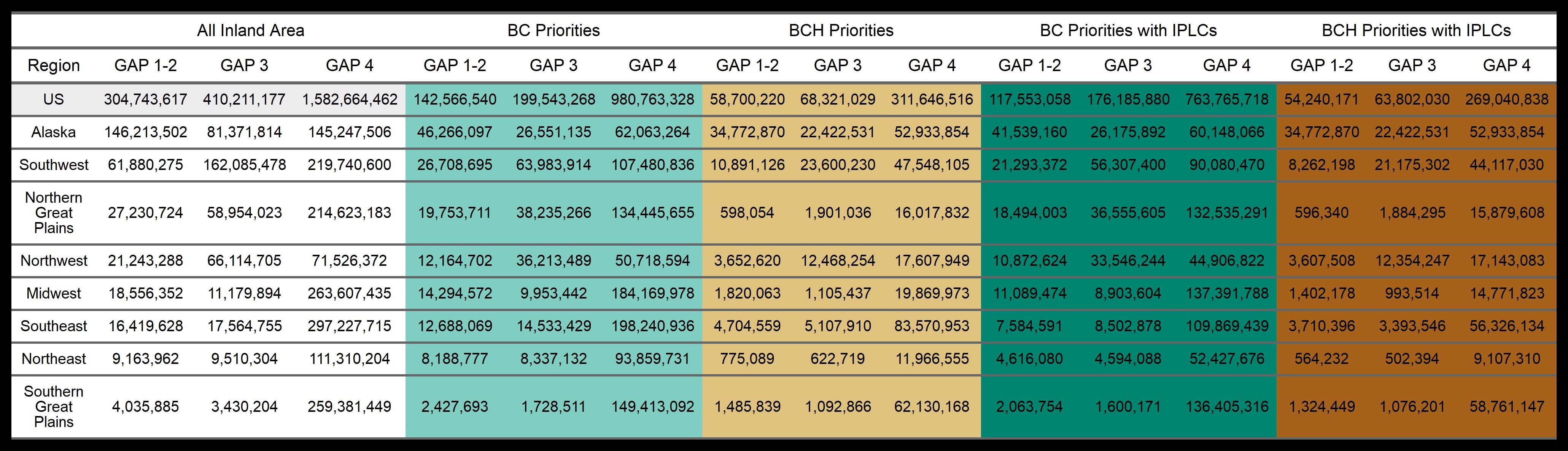

### S5 Table

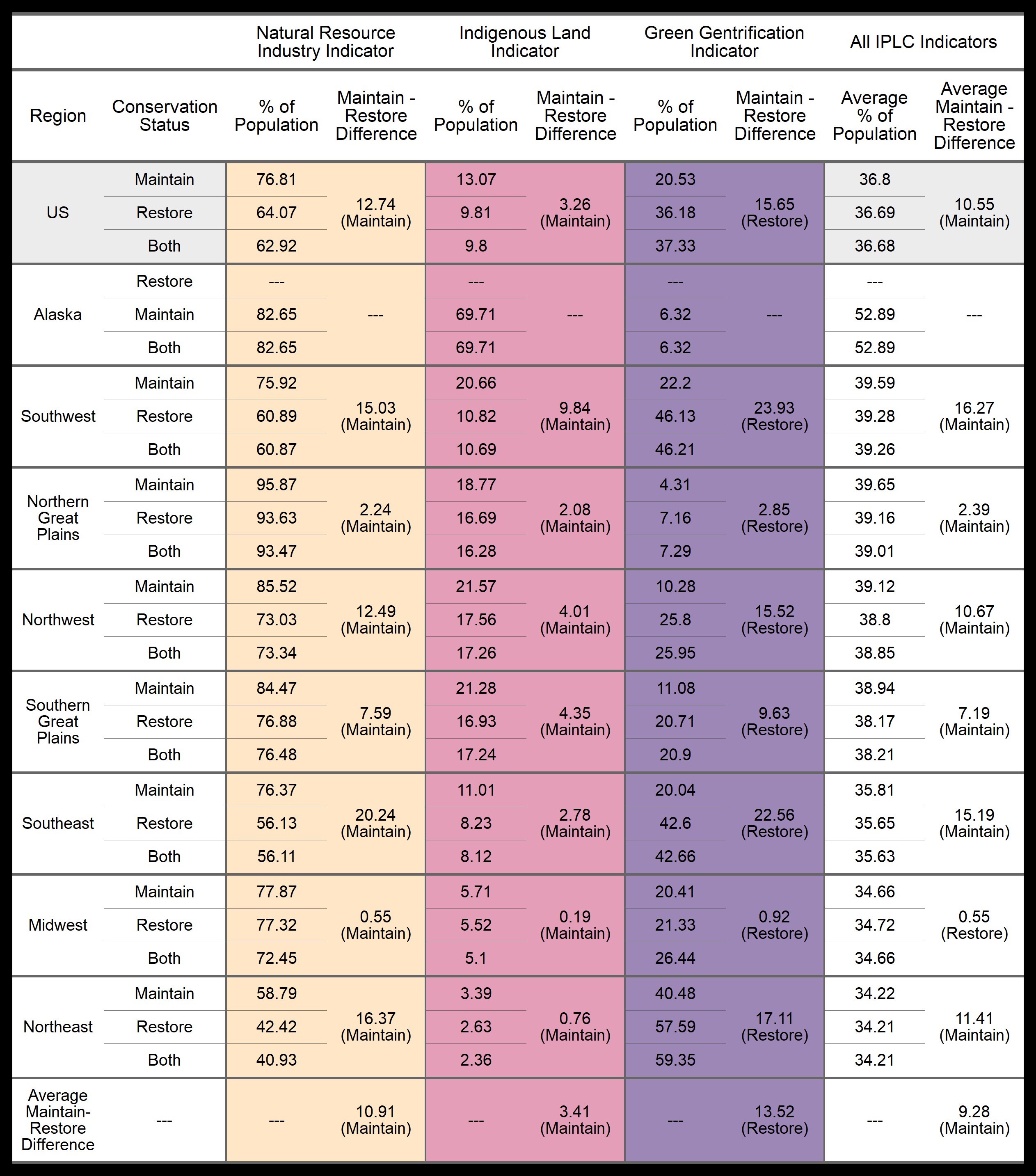
